# Amperometric reconstruction favors a toroidal fusion pore during kiss-and-run release of serotonin

**DOI:** 10.64898/2026.11.13.766543

**Authors:** Francisco F. De-Miguel

**Affiliations:** Instituto de Fisiología Celular-Neurociencias, Universidad Nacional Autónoma de México, Mexico City, Mexico

**Keywords:** kiss-and-run, fusion pore, toroidal pore, exocytosis, transmitter release, serotonin, amperometry, diffusion coefficient, hemifusion

## Abstract

Kiss-and-run amperometric currents reveal the opening and closing cycle of the fusion pore. This study uses such information along with mathematical approaches to test whether the pore is toroidal or cylindrical. Individual kiss-and-run amperometric currents from serotonin release by leech Retzius neurons were used to reconstruct either a toroid of fixed rim curvature or a cylinder of fixed length. Two pieces of information from the companion paper feed the equations: the peak pore diameter, calculated from the number of hydrated serotonin molecules released at the current peak, and the estimate of the quantal number of molecules in each event. Another essential parameter was the diffusion coefficient of serotonin, which interacts with structural parameters to determine the channel dynamics. With literature values for rim curvature (7.5 nm) and cylinder length (15 nm) and *D* value calibrated on the median event, both geometries predicted peak diameters of 2.1–3.5 nm, each requiring its own *D* (7.7% and 21.1% of the free value). Diameter scaled with resistance, with a larger exponent for the toroid (0.63 versus 0.53), indicating that toroidal pores rose more slowly and closed faster than cylindrical pores. Finding a *D* value common to both geometries left the toroid almost unchanged but shortened the cylinder to 4.1 nm, about the thickness of one bilayer. Our data favor a toroidal pore of fixed rim curvature as the structure that releases serotonin during kiss-and-run exocytosis. The short cylinder may represent the nascent pore through a hemifusion diaphragm.

**Summary:** Kiss-and-run serotonin release from leech neurons was reconstructed with toroidal and cylindrical pore models. A toroidal pore of fixed rim curvature best explains release, with a short single-bilayer cylinder as the likely nascent pore.

## Introduction

The fusion pore is the first aqueous connection between a vesicle and the extracellular space. Its geometry and dynamics determine the kinetics of transmitter release. In kiss-and-run exocytosis, the pore opens, releases part of the vesicle content, and closes again without full fusion (Staal et al., 2004; He et al., 2006; van Kempen et al., 2011; Mellander et al., 2012, 2014; Alabi and Tsien, 2013; Ren et al., 2016; Chang et al., 2022). The diameter of the open pore, and whether it flickers, closes, or dilates, determine how fast and how much of the vesicle content becomes released (Rahamimoff and Fernandez, 1997; He et al., 2006; Vardjan et al., 2007; Chowdhury and Zorec, 2020). What the pore is made of remains debated (Sørensen, 2009). Elastic models describe it as a toroidal lipid neck (Helfrich, 1973; Nanavati et al., 1992; Chizmadzhev et al., 1995, 2000). By contrast, the first conductance measurements likened it to a channel spanning two membranes (Breckenridge and Almers, 1987). Later reconstituted and cellular studies point to a pore lined by lipids and SNARE transmembrane domains, initially proteinaceous later lipidic upon dilation (Han and Jackson, 2004; Han et al., 2004; Szule and Coorssen, 2004; Zhang and Jackson, 2010; Bao et al., 2016; Sharma and Lindau, 2018).

Amperometric and conductance data alone do not resolve this question. The current fixes only the ratio between the geometric factor to the pore and the diffusion coefficient of the transmitter; conductance alone cannot distinguish a protein-lined pore from a lipidic toroid of similar size (Nanavati et al., 1992; Sharma and Lindau, 2018). Single-cell amperometry can track the expansion of individual pores (Hatami et al., 2025), but converting the current to pore size requires a model. Pore size has been estimated either by assuming a single geometry, usually a ∼15 nm long cylinder with resistivity or *D* taken from bulk solution (Wu et al., 2017), or by assuming irrelevant geometry because diffusion in the vesicle matrix limits release (Oleinick et al., 2013, 2017). In either case, features of the reconstructed pore cannot be separated from those arising from the model.

Here, we reconstruct kiss-and-run events recorded amperometrically used also in the companion paper (De-Miguel, 2026), obtained from cultured serotonergic Retzius neurons of the leech. The geometries of a toroid of fixed rim curvature and a cylinder of fixed length were compared, converting the diffusional resistance of the pore into diameter. Individual events were analyzed with their own quantal content, estimated by frozen pore extrapolation as described in the companion paper. As an independent reference, the peak diameter of the median event was matched to the hydrated flux packing (HFP) diameter of the serotonin molecules released at the current peak. Using literature values for the rim curvature and the cylinder length, each geometry required its own *D* value. Since there is no reason to suppose that each pore type operates under a different *D*, we determined a common *D* value and examined how each geometry absorbed it. These conditions supported the toroid as the pore structure, the cylinder being a candidate for the nascent pore.

## Materials and methods

Amperometric recordings of 5-HT release Amperometric currents upon 5-HT release, recorded from cultured serotonergic leech Retzius neurons, were the same as in the companion paper (De-Miguel, 2026). Pore diameters were reconstructed from these currents with the two geometries described below.

### Toroidal pore equation

Toroidal pores have no fixed length for the permeant to cross. As the pore expands or closes, the lipid membrane curves away from or toward the neck in both directions, so the bulk cylinder term is eliminated from this approximation. A toroidal neck connects two three-dimensional bulk-like regions, the vesicle lumen and the extracellular space, and material, including transmitter, converges similarly from each region into the narrow neck (Nanavati et al., 1992; Oleinick et al., 2013). The cross-sectional area of the pore, *A*(*z*), at axial position *z* is set by the minimum radius *r*_m_ and by the curvature radius of the lipid rim, *ρ*_c_. The pore radius varies as a circular arc flaring outward from the neck:

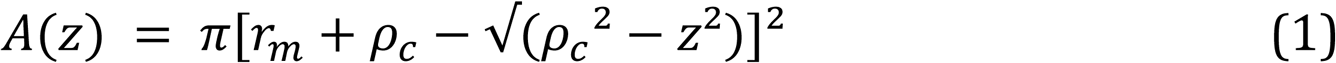

where *z* is the axial position along the neck. At *z* = 0, the narrowest cross-section, *A*(0) = π*r*_m_^2^; at *z* = ±*ρ*_c_, the profile reaches its widest point, *A*(±*ρ*_c_) = π(*r*_m_ + *ρ*_c_)^2^, where the toroid merges into the surrounding flat membrane. Because the cross-sectional area varies along the pore, its resistance is obtained by integrating the local resistivity along the axis:

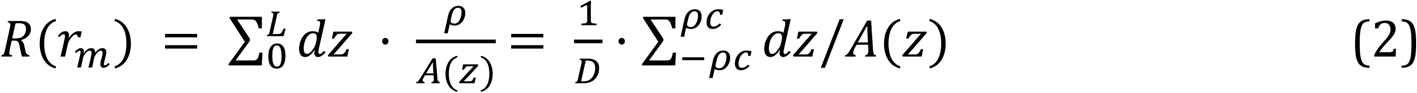

where the integral spans the axial extent of the neck, from *z* = −*ρ*_c_ to *z* = +*ρ*_c_ (*L* = 2*ρ*_c_), and *ρ* = 1/*D* is the diffusive resistivity of the medium inside the pore. The integral over the continuous flare captures in a single expression the resistances of the vesicle-facing and extracellular-facing halves of the pore. Equation 2 can be written as *R* = *L*_eff_/(*D*π*r*_m_^2^), which defines the effective length of the toroid, *L*_eff_ = π*r*_m_^2^∫d*z*/*A*(*z*), that is, the length of a cylinder of radius *r*_m_ with the same resistance. Setting the flux equal to the driving force divided by this resistance gives the implicit balance equation:

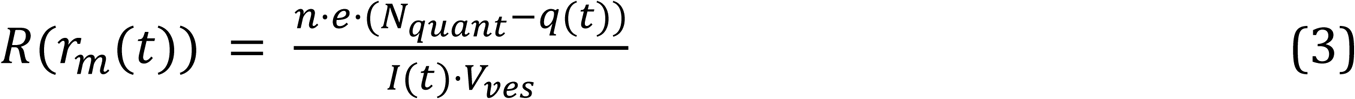

where *N*_quant_ is the quantal content of the event, estimated by frozen pore extrapolation as described in the companion paper (De-Miguel, 2026), here with the preliminary calibration of Table 1; *q*(*t*) = (1/*ne*)∫*I*(*t*′)d*t*′, integrated from 0 to *t*, is the number of molecules released up to time *t*; *n* = 4 is the number of electrons transferred per oxidized serotonin molecule (Bruns et al., 2000); *e* is the elementary charge; and *V*_ves_ is the vesicle volume. Combining Equations 1–3 gives the master equation of the toroidal pore:

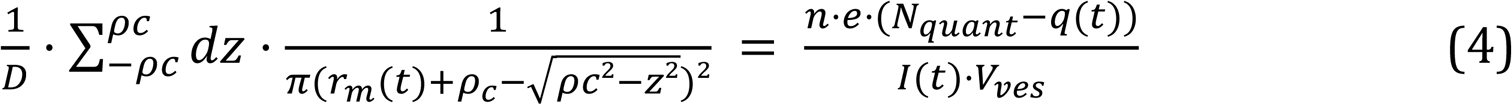

which was solved for *r*_m_(*t*) at every time point by Brent’s method, with the integral evaluated by Gauss–Legendre quadrature (80 nodes). The pore area follows directly from:

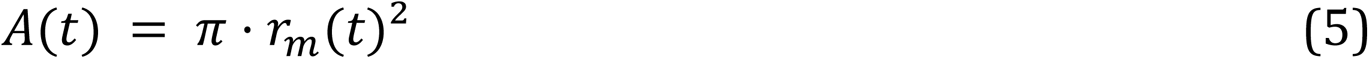

**Table 1.** Parameters of the pore reconstructions. Percentages refer to D_free_. Values marked † are derived in this work. Additional parameters, including the vesicular contents (frozen pore extrapolation), are defined in the companion paper (De-Miguel, 2026).

| Parameter | Symbol | Value | Source / note |
|---|---|---|---|
| Vesicle radius | *r*<sub>ves</sub> | 51.85 nm | Dense-core vesicles, 103.7 nm diameter (Bruns et al., 2000) |
| Vesicle volume | *V*<sub>ves</sub> | 5.84 × 10<sup>−16</sup> cm<sup>3</sup> | 4/3·π·*r*<sub>ves</sub><sup>3</sup> |
| Effective geometrical length | *L*<sub>eff</sub> | *L*<sub>Cyl</sub> + π*r*/2 (cylinder); π*r*<sup>2</sup>∫d*z*/*A*(*z*) (toroid) | Equations 2 and 7; *R* = *L*<sub>eff</sub>/(*D*π*r*<sup>2</sup>) |
| **Quantal content** |  |  |  |
| Diffusion coefficient in τ | *D*<sub>PC</sub> | 1.556 × 10<sup>−6</sup> cm<sup>2</sup>/s | Cylinder *L* = 15 nm; preliminary calibration used for the frozen pore extrapolation of the contents used in this work |
| Mean quantal size | *N*<sub>quant</sub> | 99,118 ± 4,036 | Mean ± SD (range 93,725–108,308), n = 15; preliminary frozen pore extrapolation with D<sub>PC</sub>. Final estimates: 102,650 ± 10,989 (De-Miguel, 2026) |
| **Diffusion coefficients** |  |  |  |
| Free solution | *D*<sub>free</sub> | 5.91 × 10<sup>−6</sup> cm<sup>2</sup>/s | Stokes–Einstein |
| Intravesicular | *D*<sub>ves</sub> | 2.1 × 10<sup>−8</sup> cm<sup>2</sup>/s | Marszalek et al., 1997; granule matrix in Ringer's solution |
| Cylinder | *D*<sub>Cyl</sub> | 1.25 × 10<sup>−6</sup> cm<sup>2</sup>/s (21.1%) | Calibrated on the median event † |
| Toroid | *D*<sub>Tor</sub> | 4.54 × 10<sup>−7</sup> cm<sup>2</sup>/s (7.7%) | Calibrated on the median event † |
| Common to both geometries | *D*<sub>common</sub> | 4.49 × 10<sup>−7</sup> cm<sup>2</sup>/s (7.6%) | Geometric mean of the band 3.99–5.06 × 10<sup>−7</sup> cm<sup>2</sup>/s † |
| **Pore geometry** |  |  |  |
| Cylinder length (geometry-specific D) | *L*<sub>Cyl</sub> | 15 nm | Conductance-based convention (Wu et al., 2017) |
| Toroid rim curvature radius (geometry-specific D) | ρ<sub>c</sub> | 7.5 nm | 2ρ<sub>c</sub> = L<sub>Cyl</sub>; minimum-energy pores (Nanavati et al., 1992; Jackson, 2009) |
| Cylinder length with *D*<sub>common</sub> | *L*<sub>Cyl</sub> | 4.1 nm | Median-event geometry † |
| Toroid rim curvature radius with *D*<sub>common</sub> | ρ<sub>c</sub> | 7.37 nm | Median-event geometry † |

### Cylindrical pore equation

The cylindrical pore treats the fusion pore as an ion-channel-like conductance pathway, following the cylinder-plus-access-resistance formulation of Hille (2001). It describes the outward diffusion of transmitter confined within the vesicle and the pore. The concentration of serotonin remaining in the vesicle is:

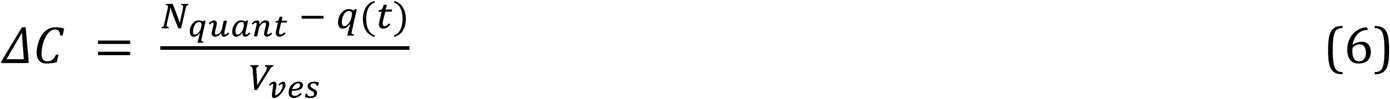

The pore is described by a length *L*_Cyl_ and a radius *r_p_*. The formulation combines a cylindrical (bulk) term with the Hall access resistance of a disk of radius *r_p_*, *R_a_* = *ρ*/(4*r_p_*) (Hall, 1975), at both the entrance and the exit of the pore; the two mouths together add *πr_p_*/2 to the length. The total resistance, produced by the bulk cylinder, *L*_Cyl_, plus the access resistance of both mouths, *πr_p_*(*t*)/2, is expressed through the effective length *L*_eff_ = *L*_Cyl_ + *πr_p_*/2:

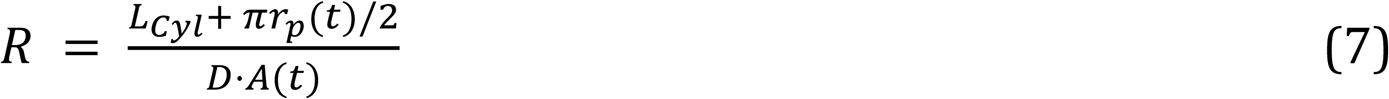

where *D* is the diffusion coefficient of serotonin within the vesicle and pore, and *A*(*t*) = *πr_p_*(*t*)^2^is the area of the pore. Equating the flux to the driving force divided by the resistance and solving for

*A*(*t*) gives the master equation of the cylindrical pore :

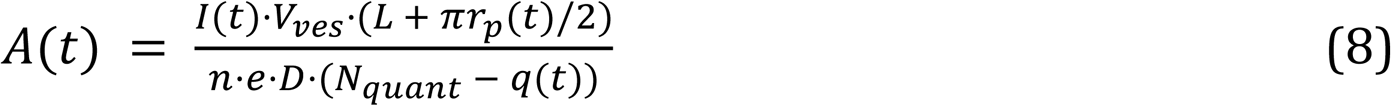

Parameters fitting the cylindrical and toroidal models With *ρ*_c_ = 7.5 nm, the semicircular neck profile (Nanavati et al., 1992) spans 2*ρ*_c_ = 15 nm along the axis, matching the length of the reference cylinder (Wu et al., 2017). This value falls within the 5.4–8.2 nm range of curvature radii for the lumen wall of minimum-energy fusion pores computed using continuum elasticity (half the separation of bilayer midplanes, 3.7–6.1 nm, plus one monolayer thickness, 1.7–2.1 nm; Jackson, 2009). The maximum pore diameter for each event was calculated as described in the companion paper (De-Miguel, 2026). A serotonin concentration of ∼270 mM, measured in Retzius neurons (Bruns et al., 2000), yields about 95,000 molecules in a dense-core vesicle 103.7 nm in diameter, close to the quantal sizes used here. The *ρ*_c_ value for the toroid and the *L*_Cyl_ value for the cylinder were obtained from the literature for *ρ*_c_ (Nanavati et al., 1992; Jackson, 2009) and for *L*_Cyl_ (Wu et al., 2017), leaving ***D*** as a free parameter. The geometry for each *D* value was calibrated on the median-amplitude event so that its peak diameter matched the center of its HFP band (5.57 molecules/µs at the current peak, rounded up to *N* = 6; *d*_H_ = 0.85 nm; 2.55 nm); the same *D* was then used for all events. Therefore, the peak diameters are predictions. Parameters and their values are listed in Table 1.

### Common diffusion coefficient

To find a *D*D value common to both geometries, the *D* required to reproduce the calibration point for three representative events (smaller, median, and larger; peak cylindrical diameter at the minimum resistance of each event) was computed as a function of the geometric parameter of each model, *D* = *L*_eff_/(*πr*^2^*R*_min_), for *ρ_c_* = 0.3–10 nm (toroid) and *L*_Cyl_ = 3–20 nm (cylinder). *D*_common_ was defined as the geometric mean of the band of *D* values across all three events in both geometries. The geometry used with *D*_common_ was the one that reproduced the median event. *D*_common_ was kept fixed for all events; peak diameters were then compared with the HFP bands of each event.

### Resistance exponent ***γ***

The dependence of throat diameter on resistance was characterized for each geometry by the local exponent:

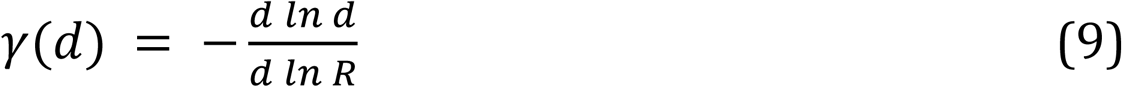

the fractional change in diameter per fractional change in resistance. *γ* was computed from Equations 2 and 7 by central differences of ln *d* against ln *R* (relative step 10^−4^). Because *R* = *L*_eff_/(*Dπr*^2^) and *D* is constant, *γ* depends only on how *L*_eff_ scales with diameter independently of *D*. Its limits are 1/2 for a long cylinder, in which *L*_Cyl_ dominates *L*_eff_ and *R* ∝ *d*^−2^; 1 for a cylinder dominated by access resistance, *R* ∝ *d*^−1^; and 2/3 for a toroid whose throat is much narrower than its rim (*r_m_* ≪ *ρ_c_*), *R* ∝ *d*^−3/2^. The characteristic exponent of each geometry was taken as the slope of ln *d* versus ln *R* between the smallest and largest peak diameters of the events (2.05–3.54 nm). To isolate the kinetic effect of *γ*, the diameter of the median event was recomputed with a constant exponent:

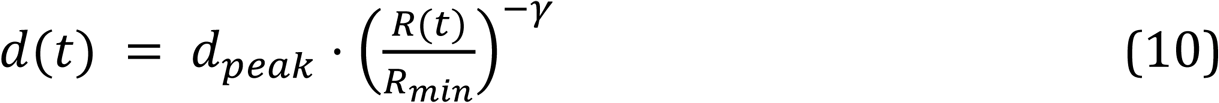

for γ = 0.5–1, keeping the measured *R*(*t*), its minimum *R*_min_ and the peak diameter *d*_peak_ (2.55 nm). Rise time was measured between 10% and 90% of the peak diameter, and half-decay time from the onset of the event to the point at which the diameter, after its peak, fell to half its peak value.

### Statistical analysis

Paired comparisons between the two reconstructions of the same events (*n* = 15) used two-sided exact Wilcoxon signed-rank tests, which make no assumption of normality; no correction for multiple comparisons was applied, and exact *p*-values are reported. The events had been recorded and classified previously (De-Miguel, 2026); neither randomization nor blinding was applicable to the reconstruction. Reconstructions, statistics and figures were computed in Python 3.11 with NumPy 2.4, SciPy 1.17 and Matplotlib 3.10. The analysis code was written with the assistance of an AI tool (Claude, Anthropic); the author checked the code and verified all results.

### Limitations

Amperometry cannot detect hemifusion, which lacks an aqueous connection; the hemifusion stage is inferred from literature, where hemifused profiles may be on-pathway or arrested intermediates and diaphragm pores remain theoretical (Zhao et al., 2016; Abbineni et al., 2018; Spencer et al., 2024). The initial pore opens abruptly to about 1 nm (Lollike et al., 1995; Zhang and Jackson, 2008), but diffusion to the electrode and filtering smooth this step and its flickers into a continuous rise (Zhang and Jackson, 2008). For a gap of 0.5–0.7 µm between the membrane and the electrode (reflecting membrane facing an absorbing electrode, *D*_free_), diffusion alone delays 90% of a step in flux by 0.4–0.9 ms. This spans the first 0.3–0.6 ms of each event, in which the reconstructed diameter falls below the hydrated diameter of serotonin (current up to 10–28% of the peak; flux ≤ 1 molecule/µs), and lowers the peak current by less than 0.5%. Sub-hydrated diameters during the rise therefore reflect a pore already open to at least one hydrated molecule, seen through the gap, rather than a smaller pore. A 1 µm gap would lower the peak by 2.5–6% and lengthen the 10–90% rise by 30–60%. The rising phase is therefore the part of the reconstruction most sensitive to the distance to the electrode, whereas the peak and the decay, which lasts tens of milliseconds, are not. In the late decay, sub-hydrated serotonin diameters appear only after 94–99% of release, at 0.2–0.4 molecules/µs, when molecules cross one at a time. Below 0.5–1 nm, transport may occur through transient free-volume elements rather than a persistent channel (Wijmans and Baker, 1995). Reconstructions are therefore most reliable near the peak (2.1–3.5 nm); because both geometries derive from the same current, this does not affect their comparison. Diffusion between the pore and the electrode adds a series resistance that does not limit the flux: an extracellular cylinder 45 nm long and 118 nm wide with *D* = 3.5 × 10^−6^ cm^2^/s contributes 0.03–0.07% of the minimum event resistance, and less as the pore closes.

Finally, because *R* = *L*_eff_/(*D*π*r*^2^), geometry and *D* cannot be separated from current, and pores this small are below the super-resolution imaging (Shin et al., 2018). Our discrimination rests on parameter plausibility, consistency across events and kinetic signatures. In nanopores of known shape, cylindrical models fit conductance less well than hourglass profiles (Kowalczyk et al., 2011), so the 4.1 nm cylinder is an effective length; the toroid approximates minimum-energy necks, which are neither exact tori nor catenoids (Nanavati et al., 1992; Jackson, 2009). Our conclusions rest on how resistance scales with throat radius rather than on the exact profile. Unlike conductance-based sizes (Wu et al., 2017), ours include the entry term, which adds about 5% to a 1 nm radius. An independent estimate of serotonin diffusivity within the pore would remove degeneracy (Yang et al., 2001).

## Results

### Reconstructed dynamics of toroidal and cylindrical kiss-and-run pores

To compare the structures of cylindrical and toroidal pores in a normalized manner using an external independent calibration, the peak diameter of the median amplitude event was matched to the mean hydrated flux packing (HFP) diameter. This was estimated from the HFP number (*N*) of serotonin molecules released at the exact current peak of each event (De-Miguel, 2026). The toroid radius of curvature (*ρ_c_*) was fixed at 7.5 nm (Nanavati et al., 1992; Jackson, 2009), while the cylinder length (*L*_Cyl_) was fixed at 15 nm (Wu et al., 2017). The diffusion coefficient *D* was the free variable, for which limited literature data are available (Renkin, 1954; Khanin et al., 1994; Marszalek et al., 1997; Dechadilok and Deen, 2006).

The reconstructed toroidal and cylindrical pores expanded similarly provided specific *D* values (Figure 1). *D*_tor_ (4.54 × 10^−7^ cm^2^/s) was 2.7 times smaller than *D*_cyl_ (1.25 × 10^−6^ cm^2^/s), yet both remained within the same order of magnitude. Referring to the theoretical free diffusion coefficient for serotonin (*D*_free_ = 5.91 × 10^−6^ cm^2^/s, estimated from the Stokes-Einstein relation), *D*_tor_ and *D*_cyl_ represented 7.7% and 21.1% of *D*_free_, respectively. The two coefficients were not interchangeable. With *D*_Tor_, the 15-nm cylinder peaked about 1.7 times higher than with its own coefficient (3.65–5.90 nm), above the HFP bands; with *D*_Cyl_, the toroid peaked at about half its own reconstruction (1.08–1.87 nm), below them (Figure 1, A″–C″).

**Figure 1.**
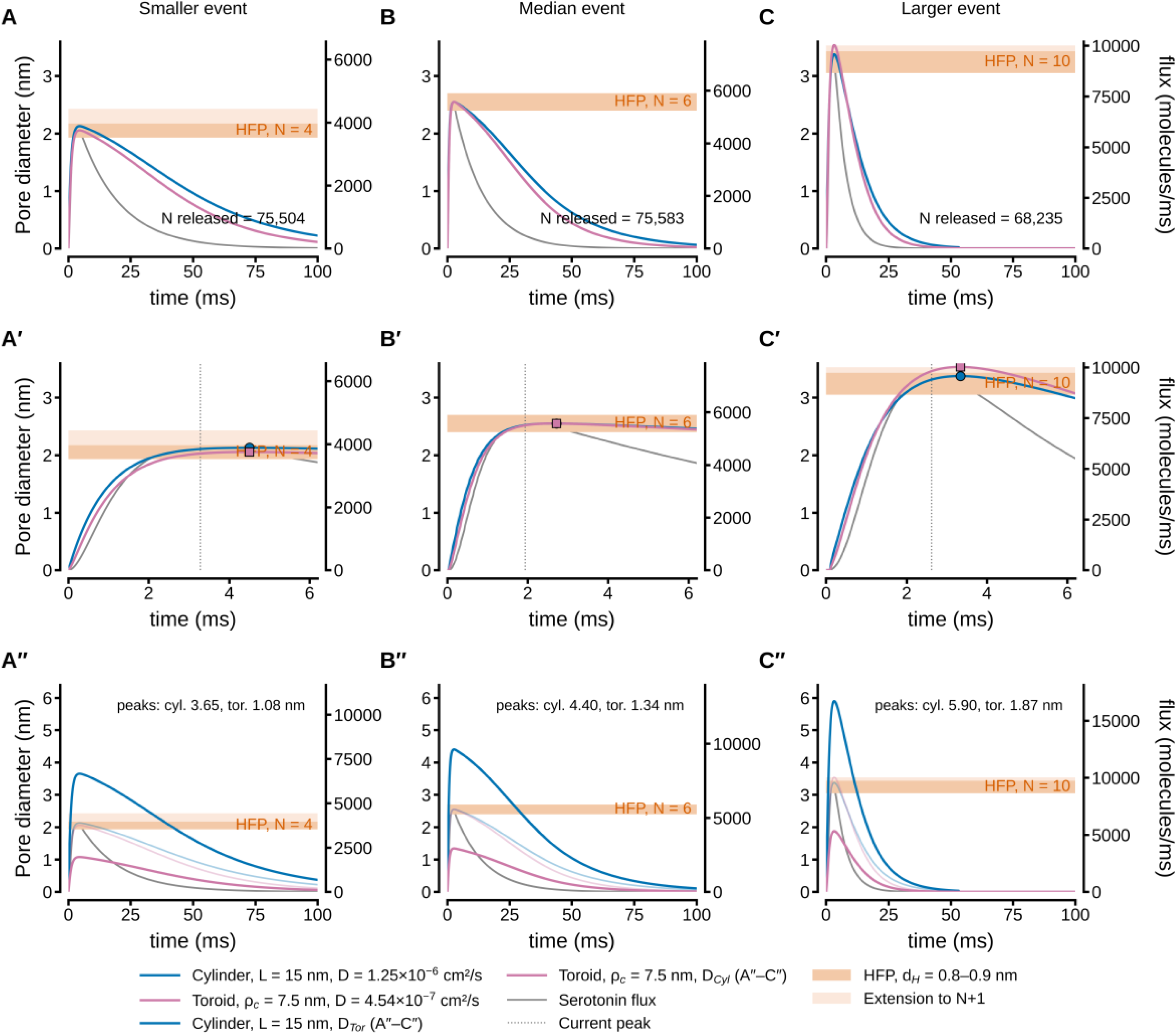
Toroidal and cylindrical reconstructions with own and exchanged diffusion coefficients. (A–C) Pore diameter (left axes) of the smaller (A), median (B), and larger (C) events, reconstructed with a cylinder (*L* = 15 nm, *D*_Cyl_ = 1.25 × 10^−6^ cm^2^/s; blue) and a toroid (*ρ*_c_ = 7.5 nm, *D*_Tor_ = 4.54 × 10^−7^ cm^2^/s; magenta). Each *D* was calibrated on the median event; peaks of the other events are predictions. Gray, measured serotonin flux (right axes), shown for timing; *N* released, number of molecules released in the event. Dark bands, HFP diameter for the *N* of each event over the range of serotonin hydrated diameter (*d*_H_ = 0.8–0.9 nm ); light bands, extension to *N* + 1. Peaks of the smaller and median events and the cylindrical peak of the larger event lie within their HFP bands; the peak of the larger toroidal event lies at the upper edge of the *N* + 1 extension (3.54 vs. 3.53 nm). (A′–C′) Rising phase of the same events. Circles and squares, cylindrical and toroidal diameter peaks (coincident in B′ by construction); dotted lines, current peak. The diameter keeps increasing for 0.7–1.2 ms after the current peak, and the two geometries diverge mainly at small diameters in the rise. (A″–C″) Same events reconstructed with the diffusion coefficient of the other geometry; faint lines, reconstructions of A–C.

A second observation was that as current amplitudes deviated from the mean, pore diameters shifted from their corresponding HFP values. Greater deviations occurred in toroidal than in cylindrical pores. A third observation was that cylindrical pores closed more slowly. When the cylindrical diameter had fallen to half its peak, the toroidal diameter was 14.3% smaller in the median event and 8.2–18.2% smaller across the fifteen events (mean 13.4%).

### The structural origin of amplitude rectification

Rectification of pore amplitudes shown in Figure 2 originates in how each geometry converts resistance into diameter. The amplitude scaling factor is the ratio *R* = *L*_eff_/(*Dπr*^2^). In the cylinder, *L*_eff_ = *L*_Cyl_ + *πr*/2, where the fixed cylinder length *L*_Cyl_ dominates the effective length. Therefore, most resistance scales with the throat area producing a resistance exponent close to 2 (*β* ≈ 1.9). In the toroid, *L_eff_* = *πr*^2^ 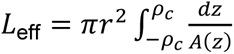, and the wall forms an arc of fixed curvature radius *ρ_c_* = 7.5 nm.

**Figure 2.**
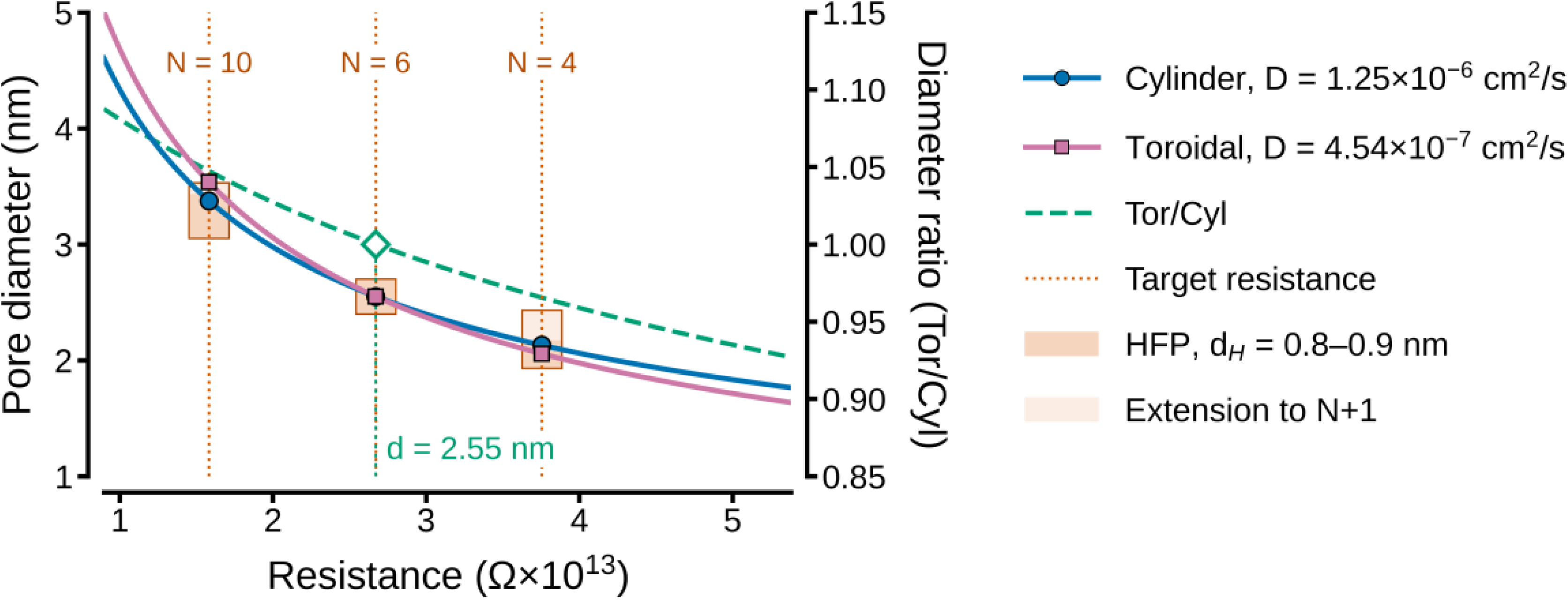
Pore diameter as a function of diffusional resistance. Cylinder (L = 15 nm, DCyl; blue) and toroid (ρc = 7.5 nm, DTor; magenta), with own calibrated D. Dotted vertical lines, minimum resistance at diameter peak of larger (N = 10), median (N = 6) and smaller (N = 4) events; circles and squares, diameters at which the cylindrical and toroidal curves reach it; boxes, HFP bands. Green dashed line (right axis), ratio of toroidal to cylindrical diameter at equal resistance; it equals 1 at d = 2.55 nm (diamond), the calibration diameter of the median event.

The local diameter rises from *d* at the throat (*z* = 0) to *d* + 2*ρ_c_* at the rim (*z* = ±*ρ_c_*) (Figure 3A). While the throat scales with *d*, the rim is set mainly by *ρ_c_*, changing little with pore size (16–19 nm for throat diameters of 1–4 nm). The flared region therefore contributes a resistance that depends weakly on *d*, resulting in a shallower overall exponent (*β* ≈ 1.6). At equal throat diameter, the toroid has about one-third of the resistance of a cylinder of the same length (*L*_eff,Tor_/*L*_eff,Cyl_ = 0.26–0.42 for *d* = 1–5 nm; 0.36 at 2.5 nm). That is why matching both geometries to the same resistance requires *D*_Tor_ ≈ 0.36 × *D*_Cyl_.

**Figure 3.**
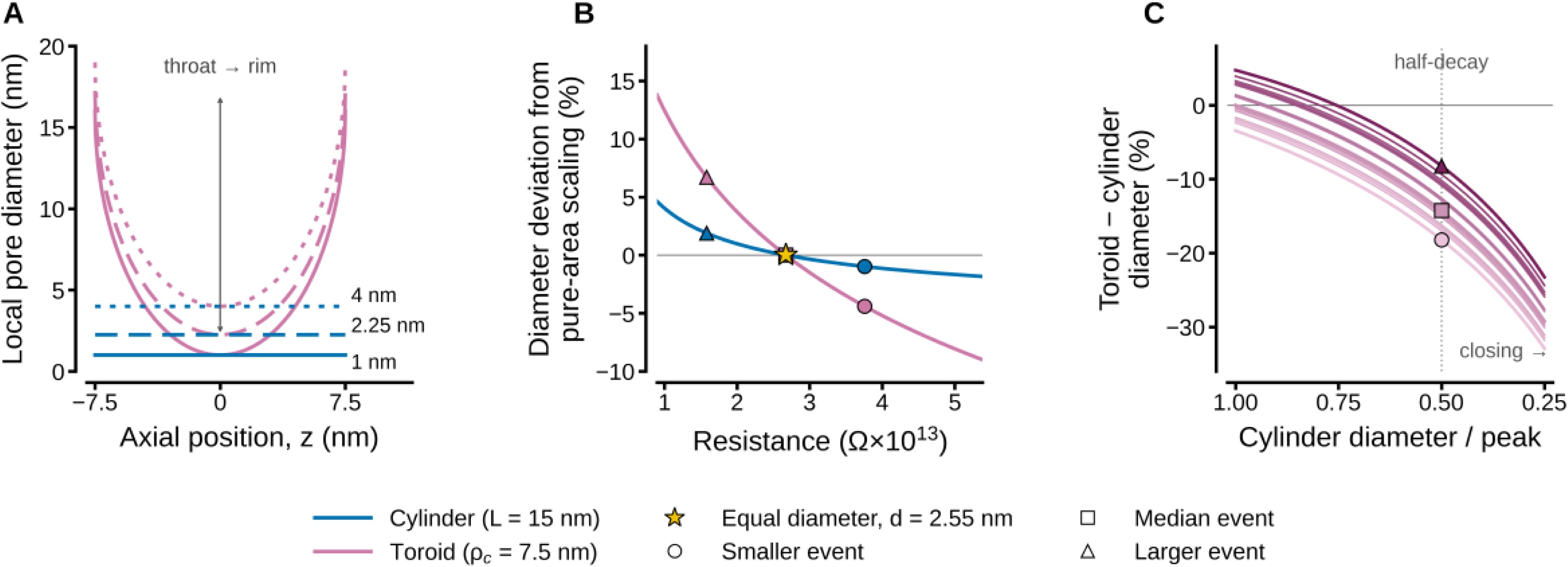
Resistance scaling explains the faster toroidal closure. (A) Local pore diameter along the axis, *z*, of toroids (*ρ_c_* = 7.5 nm; magenta) and cylinders of the same axial length (*L* = 15 nm; blue) at throat diameters of 1, 2.25, and 4 nm. The toroidal throat follows *d*, whereas the rim diameter (*d* + 2*ρ_c_* = 16–19 nm) is set mainly by *ρ_c_*. (B) Deviation of the diameter from pure-area scaling (*d* ∝ *R*^−1/2^), normalized to the resistance at which both geometries give the same diameter (star; *d* = 2.55 nm, median event). Both deviations are negative above that resistance and positive below it; the toroidal deviation is three to five times the cylindrical one. Circles and triangles represent peak resistances of the smaller and larger events, respectively. (C) Toroid–cylinder diameter difference during closure, expressed as a percentage of the cylindrical diameter, plotted against the cylindrical diameter relative to its peak (from 1.00 down to 0.25). Fifteen events are shown, ordered from smaller (light) to larger (dark); circle, square, and triangle indicate the smaller, median, and larger events at half-decay (dotted line), where toroidal diameters are 8.2–18.2% smaller than cylindrical ones.

### An exponent shapes the rise and decay of pore diameter

The toroidal pore rises more slowly than the cylindrical pore and closes faster (Figure 1A′–C′; Figure 3C). Both features have the same origin (Equation 9). The diameter depends on resistance according to *d* ∝ *R*^−*γ*^ . Because both reconstructions are monotonic transforms of the same *R*(*t*), differences in the shape of their time courses arise only from the effective length of each geometry, *L*_eff_(*r*) (Equations 2 and 7); *D* and the constants *n*, *e*, and *V*_ves_ scale the amplitude but not the shape. For narrow throats, *ρ_c_* is purely an amplitude factor in *L*_eff_/*D*. Over the diameters of the events, *γ* = 0.63 for the toroid, 0.53 for the cylinder, and 0.5 for pure-area scaling. Therefore, the toroid departed further from pure-area scaling over resistances of 0.9–5.4 × 10^13^ Ω, and normalized where both geometries coincided, −9.0% to +14%, compared with −1.8% to +4.6% for the cylinder. At the peaks of the smaller and larger events, the departures were −4.4% and +6.7% for the toroid and −1.0% and +1.9% for the cylinder (Figure 3B).

Consequently, the diameter ratio *d*_Tor_/*d*_Cyl_ decreases monotonically from 1.09 to 0.93 as resistance rises, crossing 1 at the median event (2.67 × 10^13^ Ω, *d* = 2.55 nm), where both *D* values were calibrated. Because *R* falls during opening and rises again during closure, the same exponent acts in opposite directions in the two phases. To reach the same peak, the toroidal throat changed diameter ∼ 20% faster than the mouth of the cylinder (*γ*_Tor_/*γ*_Cyl_ ≈ 1.2).

Peak diameter and minimal resistance coincide in both geometries (2.7 ms in the median event). However, the toroid is delayed in reaching 10% of its peak (0.04–0.08 ms), and in reaching a 0.65 nm diameter (0.06–0.12 ms). Its 10–90% rise time is 3.9% and 2.5% longer in the smaller and median events, and equal in the larger event (−0.3% ; median +1.1% ). At the peak, both extreme size reconstructions differed from −3.5% (smaller) to +4.7% (larger) and agreed with the predicted HFP values. For the larger event, the cylindrical peak (3.38 nm) layed within its *N* = 10 band; the toroidal peak (3.54 nm) layed at the upper edge of the *N* + 1 extension (3.53 nm). For the smallest event, both peaks layed within its N = 4 band. During closure, the toroid shrank faster and divergence grew as pores narrowed. At half-decay the toroid was 8.2–18.2% smaller than the cylinder, and at a quarter of the peak 23–33% smaller. The largest differences occurred in events with smaller peaks. Those events started closer to the small-diameter region. These differences depended on the calibration. Each geometry was matched to the HFP diameter of the median event with its own *D* (*D*_Tor_ ≈ 0.36*D*_Cyl_ ). For this reason, the crossing of *d*_Tor_/*d*_Cyl_ at the median event, the rectification of peak diameters, and part of the kinetic contrast reflected the *D* pair as much as the pore shape. Because *R* = *L*_eff_/(*D*π*r*^2^), *D* traded off against the geometric parameters *L*_Cyl_ and *ρ_c_*. Therefore, a toroid and a cylinder reconstructed with different coefficients cannot be compared on equal terms. Separating the contribution of pore shape from that of diffusion required a single *D* for both geometries. The following sections examine how the exponent shaped the kinetics, how *D* traded off against *ρ_c_* and *L*_Cyl_, and whether a common *D* reproduced the HFP calibration in both geometries.

### ***γ*** and the kinetics of pore diameter

*γ* provides relevant information for condensing the geometrical information independent of *D*. To isolate the effect of *γ*, we used Equation 10, *d*(*t*) = *d*_peak_ ⋅ (*R*(*t*)/*R*_min_)^−*γ*^for the median event, maintaining identical *R*(*t*) and the peak diameter profiles (Figure 4C–E). Increasing *γ* from 0.5 to 1.0 lengthened the 10–90% rise time from 1.04 to 1.15 ms and shortened the half-decay time from 33.7 to 22.8 ms without shifting the time to peak. At the characteristic exponents of the two reference geometries this single-parameter approximation closely reproduced the full reconstructions. For the cylinder *γ* = 0.53 it yielded a rise time of 1.05 ms and a half-decay time of 32.7 ms (compared to 1.06 ms and 32.9 ms for the full model); for the toroid *γ* = 0.63 yielded a rise time of 1.09 ms and a half-decay time of 29.6 ms (compared to 1.09 ms and 29.4 ms). Although at first glance the differences seem small, the kinetic impact of *γ* is notably asymmetric. Shifting from 0.53 in cylinder to 0.63 in toroidal lengthens the rise time by 3.5% but shortens the half-decay time by 9.5%. Because γ affects the decay nearly three times more than the rise, closure is the most robust kinetic difference between the two geometries, whereas rise time is the most sensitive.

**Figure 4.**
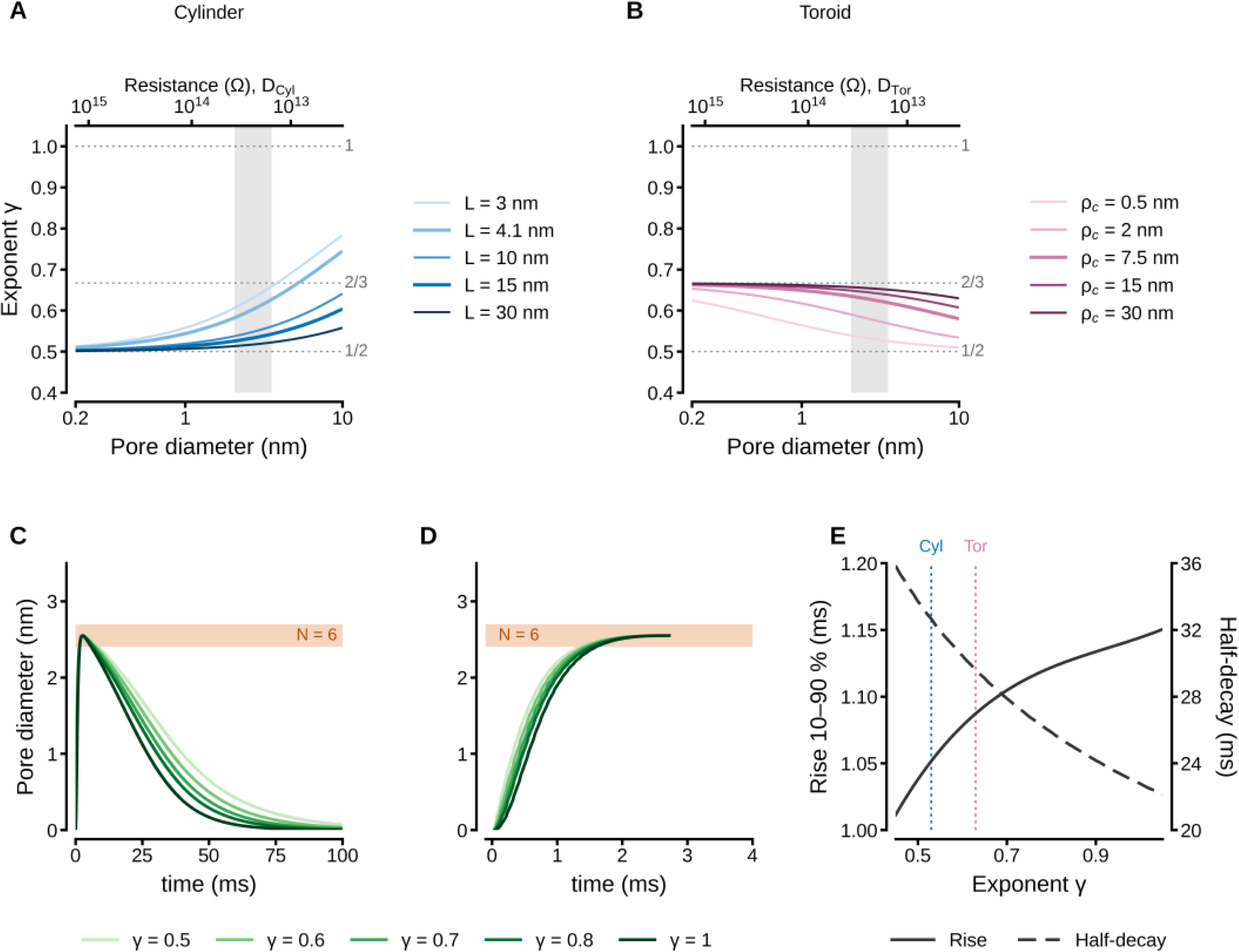
The local exponent *γ* determines the shape of the reconstructed pore dynamics. (A, B) Exponent *γ* = −*d*(*ln d*)/*d*(*ln R*) versus pore diameter for cylinders with *L*_cyl_ = 3–30 nm (A) and toroids with *ρ_c_* = 0.5–30 nm (B). Thick lines indicate reference geometries (*L*_cyl_ = 15 nm; *ρ_c_* = 7.5 nm ) and the common-*D* cylinder (*L*_cyl_ = 4.1 nm ); the common-*D* toroid (*ρ_c_* = 7.37 nm ) is indistinguishable from the reference toroid. Gray band highlights the range of event diameters (2.1–3.5 nm); dotted lines mark asymptotes at 1/2, 2/3, and 1. Top axes show the resistance of each reference geometry with its own *D*, provided as a guide. (C, D) Diameter of the median event computed as *d*(*t*) = *d*_peak_ ⋅ [*R*(*t*)/*R*_min_]^−*γ*^for *γ* = 0.5–1, showing the full time course (C) and the rising phase (D). Bands represent the HFP diameter for *N* = 6 (dark) and extension to *N* + 1 (light). (E) 10–90% rise time (solid line, cubic trend) and half-decay time (dashed line) versus *γ*; vertical lines mark the reference cylinder (0.53) and toroid (0.63).

### Trade-off between geometry and D

Because 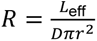, any variation in *D* can be accounted for by adjusting the geometric parameters of each model. Using *D*_Tor_ = 4.54 × 10^−7^ cm^2^/s, varying the toroidal rim curvature from *ρ*_c_ = 0.3 nm (the smallest radius of a lipid pore resolved in atomistic simulations; Wohlert et al., 2006) to 7.5 nm (Jackson, 2009) shifted the diameter that matches the HFP reference by ≈ 1.9 nm (0.69–2.55 nm; Figure 5A). By contrast, using *D*_Cyl_ = 1.25 × 10^−6^cm^2^/s and varying the cylinder length from *L*_Cyl_ = 5 to 20 nm shifted it by ≈ 1.4 nm (1.54–2.92 nm; Figure 5C). Either parameter moved the calibration point across the whole range of kiss-and-run diameters, with an effect of the same order as changing *D*. However, the two geometries differ in the cost of that freedom. Imposing *D*_Cyl_ on the toroid required *ρ*_c_ = 47.7 nm (Figure 5B), close to the 51.85 nm vesicle radius during full fusion; imposing *D*_Tor_ on the cylinder required *L*_Cyl_ = 4.2 nm (3.6 times shorter than 15 nm; Figure 5D). This length is compatible with the 4–5 nm thickness of a single lipid bilayer. In either case, the alternative and the reference curves remained nearly superimposed over the relevant range of diameters, so current alone cannot resolve either degeneracy. Only the physical admissibility of parameters discriminates between them.

**Figure 5.**
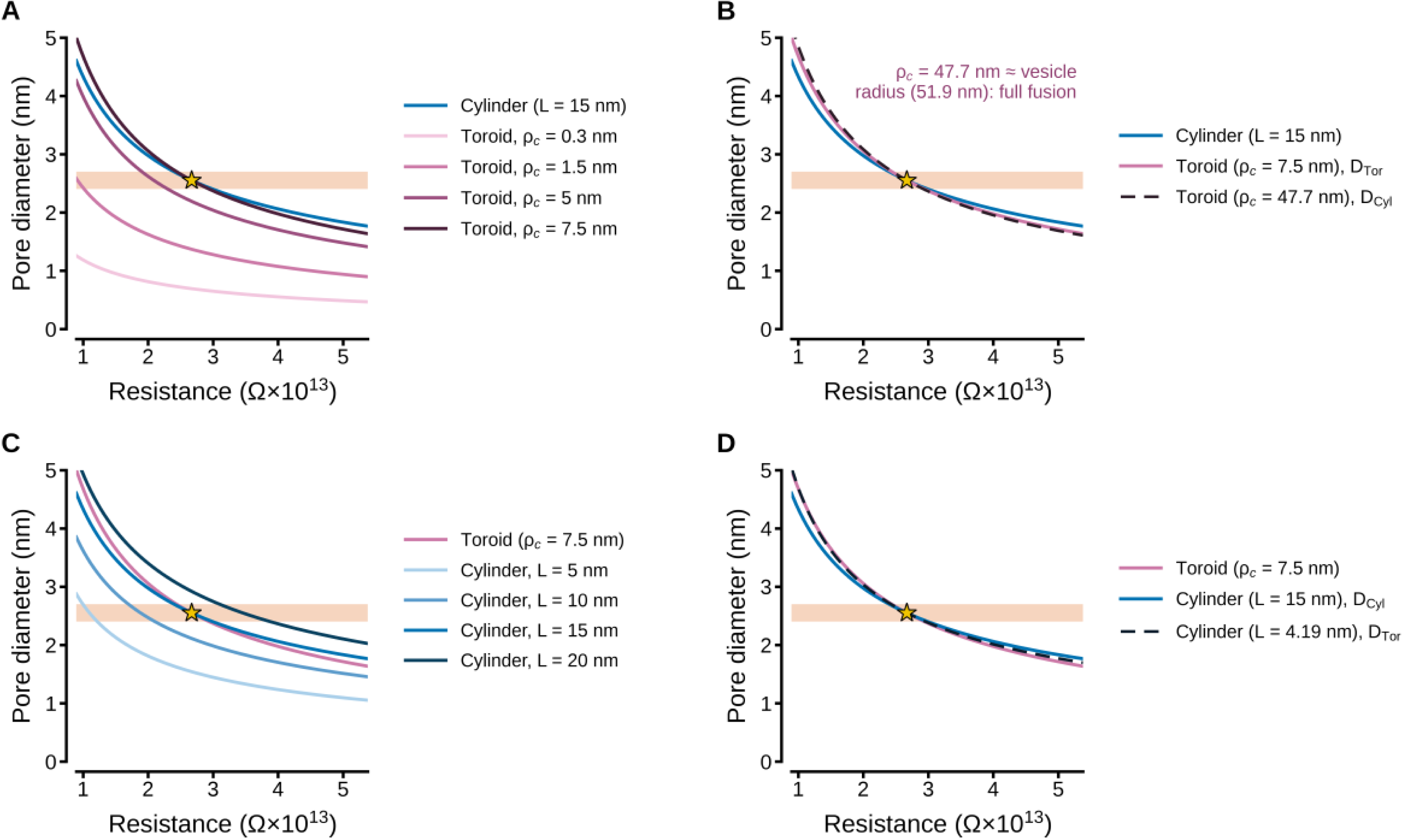
Trade-off between geometric parameters and diffusion coefficient. Pore diameter as a function of resistance. Star indicates the calibration point of the median event (d = 2.55 nm); dark band, HFP diameter for N = 6 ; light band, extension to N + 1 . (A) Toroids with ρ_c_ = 0.3–7.5 nm, all using D_tor_, alongside the reference cylinder (*L_Cyl_* = 15 nm, D_cyl_ ). (B) Reference cylinder and reference toroid, each evaluated with its own D, and the toroid that reproduces the calibration point using D_cyl_ (dashed line), which requires ρ_c_ = 47.7 nm (roughly the vesicle radius). (C) Cylinders with L = 5–20 nm, all using D_cyl_, alongside the reference toroid (ρ_c_ = 7.5 nm, D_tor_). (D) Reference toroid and reference cylinder, each evaluated with its own D, and the cylinder that reproduces the calibration point using D_tor_ (dashed line), which requires L = 4.19 nm.

Extending the analysis to a continuous range of *D* showed how each geometry absorbs a change in the *L*_eff_/*D* coefficient. For the cylinder, the effective length that reproduced the calibration point was a linear function of *D*. Ths was because *L*_Cyl_ enters additively in *L*_eff_ = *L*_Cyl_ + *πr*/2 . It rises from 0 nm at *D* = 1.5 × 10^−7^ cm^2^/s (2.5% of *D*_free_) to 79 nm at *D*_free_; the 3–20 nm range corresponds to 6–27% of *D*_free_ (Figure 6). For the toroid, *ρ_c_* grew superlinearly with *D* and reached the 51.85 nm vesicle radius at *D* = 1.30 × 10^−6^cm^2^/s, about 22% of *D*_free_; the toroid is admissible only well below this value, a range that contains *D*_Tor_ (7.7%) and *D*_common_ (7.6%) but not *D*_Cyl_ (21.1%). The two dependencies propagated differently. In the cylinder, an uncertainty in *L* translated into a proportional uncertainty in *D* over the whole range, whereas in the toroid the same absolute uncertainty in *ρ_c_* changed *D* strongly when the rim was tight and weakly near *ρ_c_* = 7.5 nm.

**Figure 6.**
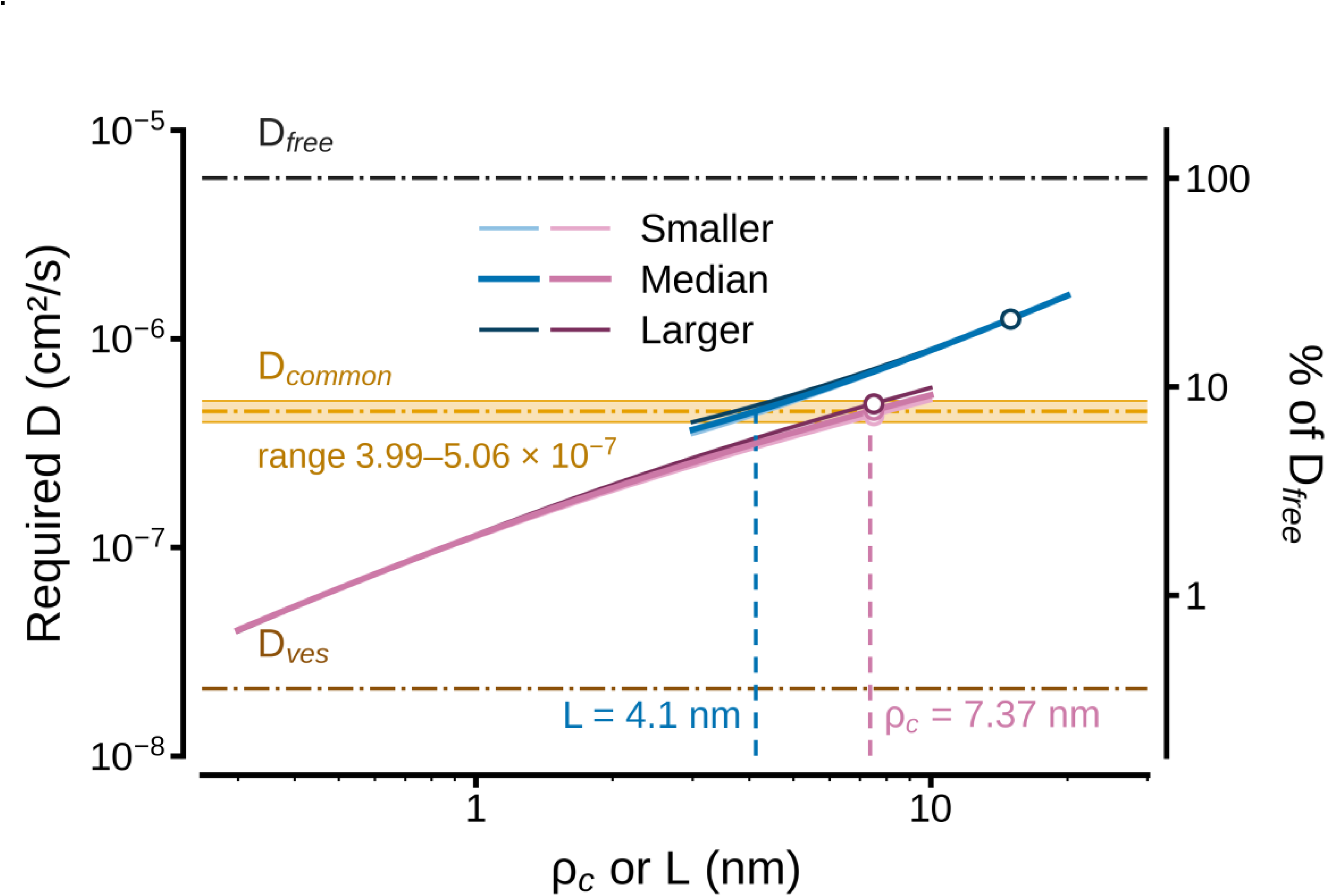
Diffusion coefficient required to reproduce the HFP calibration. Required D as a function of ρ_c_ for the toroid (magenta) and of L_Cyl_ for the cylinder (blue), for the smaller (light), median (medium) and larger (dark) events, each with its own N_quant_. Circles, reference geometries (ρ_c_ = 7.5 nm; L_Cyl_ = 15 nm). Orange band, D_common_ range (3.99–5.06 × 10^−7^ cm^2^/s); orange dash-dot line, D_common_ = 4.49 × 10^−7^ cm^2^/s. Upper and lower dash-dot lines, D_free_ and D_ves_. Vertical dashed lines, L_Cyl_ = 4.1 nm and ρc = 7.37 nm, the geometry at which the median event reaches D_common_. Right axis, D as a percentage of D_free_.

### A common diffusion coefficient for both geometries

Because *D* is a pure scaling factor, the coefficient that reproduces the HFP diameter can be solved at any fixed geometry. Figure 6 shows this coefficient as a function of *ρ_c_* and *L*_Cyl_ for the smaller, median, and larger events. For both geometries, the required *D* increased with the geometric parameter. This happens because a wider rim or a longer channel raises *L*_eff_ at a given diameter. The curves of the three events nearly coincide. At the reference geometry, the toroid required 4.30 × 10^−7^ cm^2^/s, 4.54 × 10^−7^ cm^2^/s, and 4.90 × 10^−7^ cm^2^/s (smaller, median, and larger events), whereas the three cylinder curves coincided at 1.25 × 10^−6^ cm^2^/s because the target diameters were the cylinder peaks. The toroidal and cylindrical curves meet only in a narrow region near 4.5 × 10^−7^ cm^2^/s, where the toroid remained close to its reference rim curvature and the cylinder was at the short end of its range.

The band was bounded by the explored parameter ranges. Its lower limit was set by the cylinder of the larger event at *L*_Cyl_ = 3 nm and its upper limit by the toroid of the smaller event at *ρ*_c_ = 10 nm, so *D*_common_ is an operational value defined by the overlap of the curves within those ranges. Taken as the geometric mean of the band, *D*_common_ = 4.49 × 10^−7^ cm^2^/s (band 3.99–5.06 × 10^−7^ cm^2^/s; 7.6% of *D*_free_ ) lay within 1.1% of *D*_Tor_ (4.54 × 10^−7^ cm^2^/s). In practice, a common coefficient amounts to imposing the toroidal *D* on the cylinder, which accordingly shortened to 4.1 nm. That is practically the 4.2 nm obtained above with *D*_Tor_.

### Diameter dynamics with fixed median geometry

Because the current fixes only the ratio *L*_eff_/*D*, a common coefficient was possible only if the *L*_eff_ of each model changes to preserve that ratio. With *D*_common_ (64% smaller than *D*_Cyl_ ) and the geometry that reproduces the median event (Figure 7), the toroidal rim curvature changed by only 2%, from 7.5 to 7.37 nm, whereas the cylinder shortened by 72%, from 15 to 4.1 nm, a length that covers only the throat of the toroidal profile (Figure 8A). The differences between the two frameworks therefore arose almost entirely from the cylinder.

**Figure 7.**
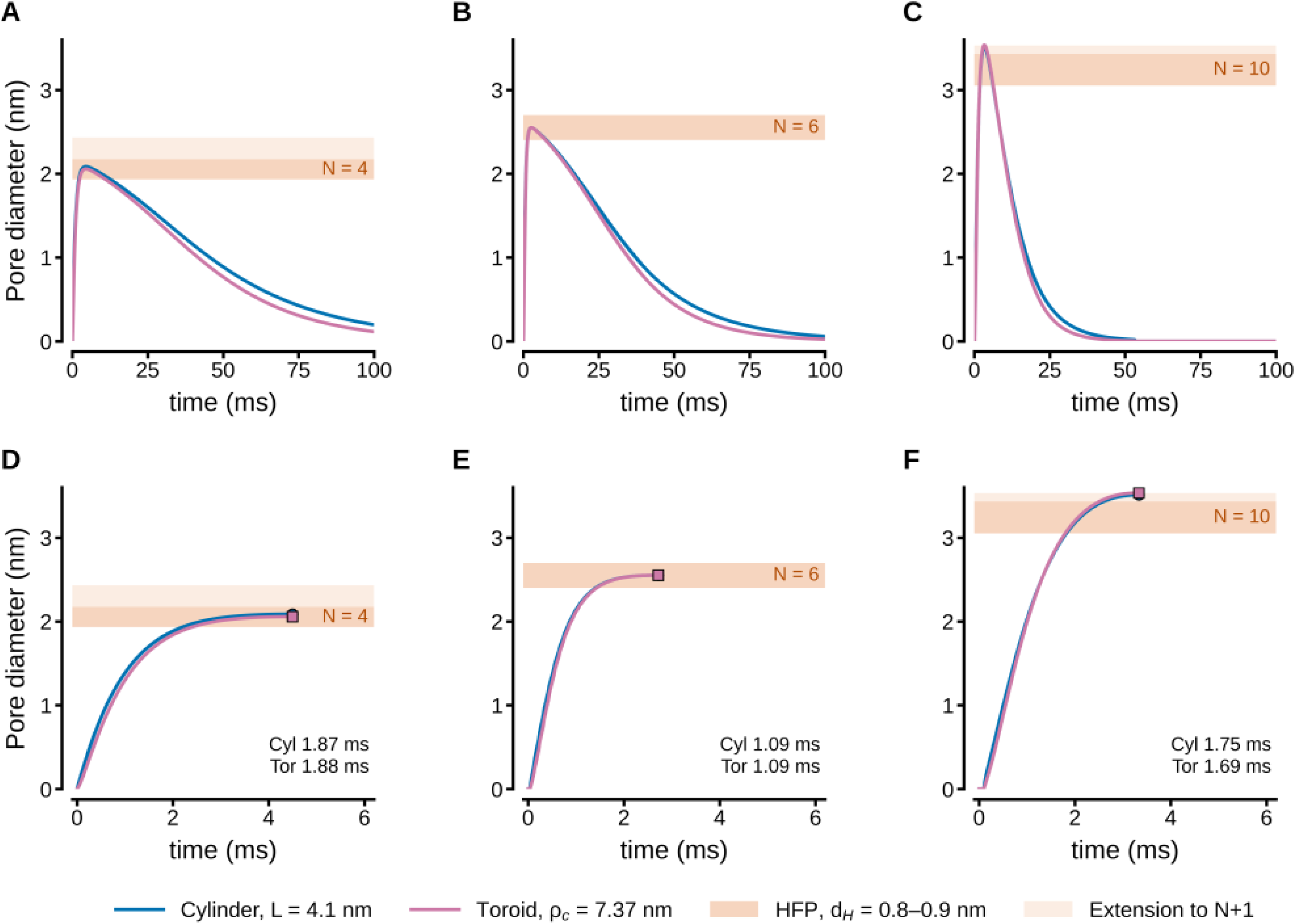
Pore diameter with a common diffusion coefficient. (A–C) Full time course and (D–F) rising phase of the smaller (A, D), median (B, E), and larger (C, F) events, reconstructed with *D*_common_ = 4.5 × 10^−7^ cm^2^/s and the median-event geometry, fixed for all events: cylinder, *L*_Cyl_ = 4.1 nm (blue); toroid, *ρ_c_* = 7.4 nm (magenta). Bands are defined as in Figure 1. Circles and squares indicate cylindrical and toroidal diameter peaks, respectively; numbers show 10–90% rise times.

**Figure 8.**
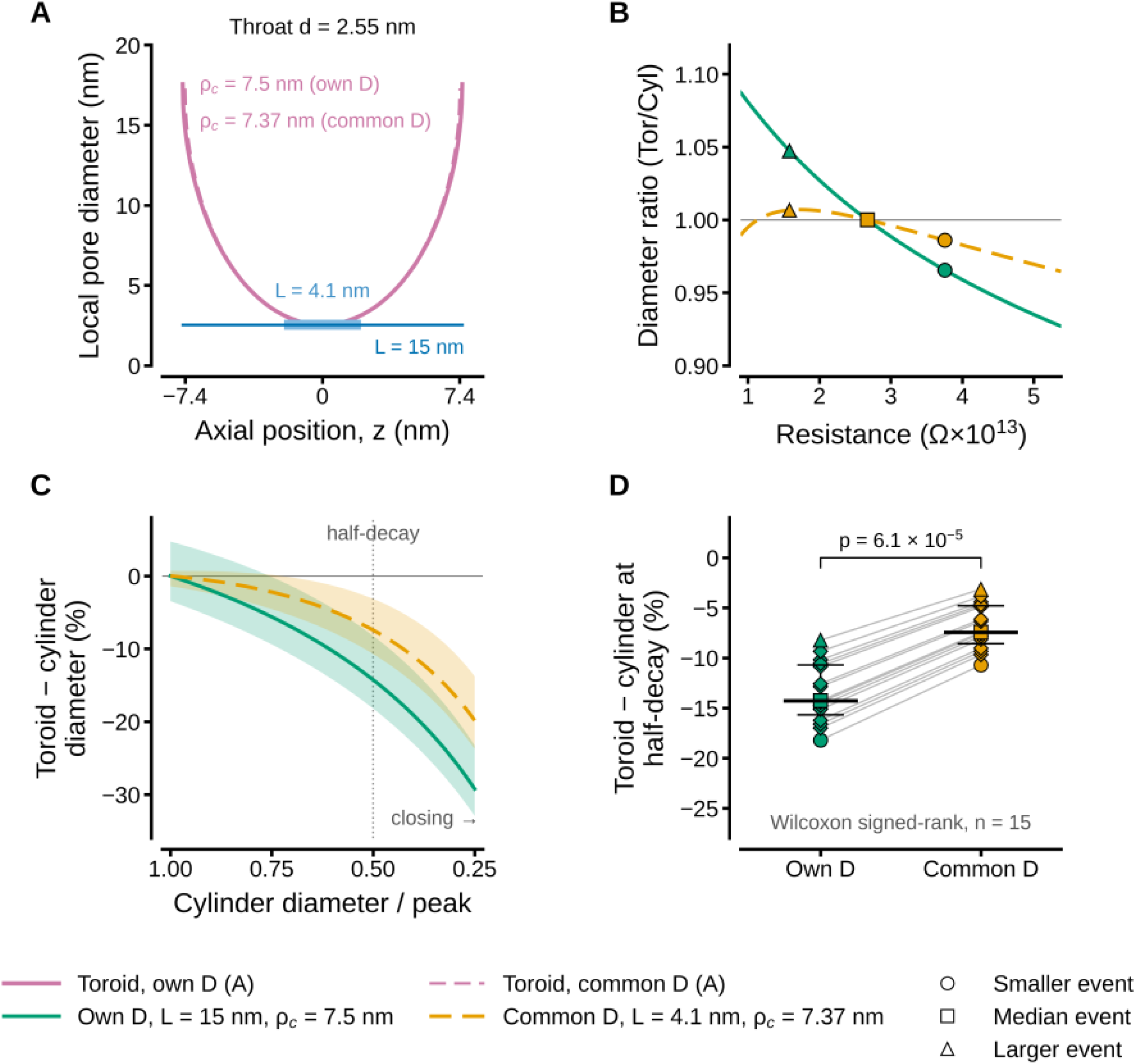
A common diffusion coefficient reduces the contrast between toroidal and cylindrical reconstructions. (A) Wall profiles at a throat diameter of 2.55 nm (calibration diameter of the median event): toroid evaluated with its own *D* (*ρ_c_* = 7.5 nm; solid magenta line) and with *D*_common_ (*ρ_c_* = 7.37 nm; dashed magenta line), alongside cylinders of 15 nm (blue) and 4.1 nm (light blue) length. (B–D) Green indicates geometry-specific *D* (*L*_Cyl_ = 15 nm, *ρ_c_* = 7.5 nm); orange indicates *D*_common_ (*L*_Cyl_ = 4.1 nm, *ρ_c_* = 7.37 nm). (B) Ratio of toroidal to cylindrical diameter (*d*_tor_/*d*_cyl_) at equal resistance; circle, square, and triangle indicate peak resistances of the smaller, median, and larger events, respectively. (C) Toroid–cylinder diameter difference during closure as a function of the cylindrical diameter relative to its peak: line shows the median of fifteen events, and shaded band represents the range; dotted line marks half-decay. (D) Toroid–cylinder diameter difference at half-decay for each of the fifteen events; gray lines connect values from the same event; thick horizontal bars denote medians, and thin bars denote interquartile ranges (*p* = 6.1 × 10^−5^, two-sided exact Wilcoxon signed-rank test, *n* = 15).

Imposing this geometry on all events reproduced the median event by construction and placed every other event within its HFP band or its *N* + 1 extension. The larger event peaked at 3.51 nm (cylinder) and 3.53 nm (toroid), at the upper edge of the *N* + 1 extension of its *N* = 10 band (3.53 nm), with a cylinder peak 4.0% larger than with geometry-specific coefficients (3.38 nm).

Rectification of peak diameters kept its direction. The toroid remained smaller than the cylinder in small pores and larger in large pores; its magnitude decreased. The *d*_Tor_/*d*_Cyl_ ratio of the smaller and larger events changed from 0.97 and 1.05 to 0.99 and 1.01 (Figure 8B). The ratio of the median event remained 1 by construction. Across all events, the deviation of the ratio from unity at the peak decreased from a median of 2.0% to 0.6% (two-sided exact Wilcoxon signed-rank test, *n* = 15, *p* = 1.2 × 10^−4^ ). With *D*_common_, the ratio peaked at 1.007 near 1.7 × 10^13^ Ω, close to the resistance of the larger event (1.58 × 10^13^ Ω), and decreased at lower resistances (Figure 8B).

The magnitudes of rectification and of the kinetic asymmetry are set by how each *L*_eff_ scales with diameter. In the cylinder, *γ* rises from 0.5, where the channel term dominates, toward 1, where the access term dominates; a shorter channel reaches the access regime at smaller diameters (Figure 4A). In the toroid, the rim acts mainly as an amplitude factor interchangeable with *D*. Therefore, *γ* stays ∼2/3 and is almost insensitive to *ρ*_c_ (Figure 4B). Over the diameters of the events, the *D*_common_ geometry therefore moves the cylinder toward the toroid: shortening it from 15 to 4.1 nm increased its exponent from 0.53 to 0.60, whereas the toroidal exponent remained at 0.63. The Tor/Cyl exponent ratio dropped from 1.18 to 1.05.

The kinetic consequences followed this ratio. Both geometries yielded nearly identical peak diameters in both frameworks (*p* = 0.36), and the toroid still closed faster, but by less. At half-decay, the toroidal pore had a diameter 14.3% smaller than the cylinder with geometry-specific coefficients, compared to 7.4% smaller with *D*_common_ (medians; smaller in all fifteen events, *p* = 6.1 × 10^−5^; Figure 8C, D). The rise asymmetry, which was smaller, reversed its sign. The toroid grew more slowly than the cylinder with geometry-specific coefficients (median +1.1%) but faster in fourteen of fifteen events with *D*_common_ (median −1.9%; *p* = 6.1 × 10^−5^).

The toroidal half-decay time remained shorter in all events in both frameworks, by a median of 10.6% (range 7.5–15.9%) with geometry-specific coefficients and 5.7% (range 2.7–7.7%) with *D*_common_ (*p* = 6.1 × 10^−5^). The toroidal times barely changed between frameworks (rise 1.9, 1.1, and 1.7 ms; half-decay 40.7, 29.4, and 12.0 ms for the smaller, median, and larger events), so the reduction of the asymmetry comes from the cylinder, which, shortened to 4.1 nm, rises more slowly and closes faster.

As expected, both reconstructions peaked at the same instant, since both are monotonic transforms of the same *R*(*t*) (therefore *I*(*t*)). The rise asymmetry, however, reversed sign between frameworks, whereas faster toroidal closure persisted. This is therefore the only kinetic signature independent of the choice of *D* . Adopting a common coefficient transferred the geometric discrepancy from *D* to *L*_Cyl_ . A cylinder of single-bilayer length brings its diameter–resistance relation close to that of the toroid. Conversely, the differences obtained with geometry-specific coefficients reflect the contrast between a pore whose curved rim does not scale with its throat and a channel spanning two apposed membranes.

Why a common coefficient requires a cylinder about 4 nm long stems from the resistances of the two geometries at equal *D*. For a given rim curvature radius, the ratio L_eff,Tor_/L_eff,Cyl_ has a maximum at an intermediate throat diameter. In addition, the two geometries can reach the same resistance at some diameter only if the cylinder is shorter than a critical length at which this maximum equals one. Numerically, *L*_crit_ = 0.57*ρ*_c_ for *ρ*_c_ = 0.5–10 nm (Figure 9A). For *ρ*_c_ = 7.37 nm, *L*_crit_ = 4.21 nm, with the tangency at *d* = 3.39 nm, within the diameters of the events. The common-*D* cylinder (4.1 nm) lies just below this limit (Figure 9B). Its resistance equals that of the toroid at the calibration diameter (2.55 nm) and stays within 2.5% of it in all events (2.05–3.54 nm). This explains both the length of the common-*D* cylinder and the near coincidence of the two reconstructions with *D*_common_ (Figure 8B). Conversely, the reference cylinder (15 nm) lies far above *L*_crit_, and its resistance exceeds that of the toroid at every diameter *(L_eff,Tor_/L_eff,Cyl_ = 0.34–0.39)*, which is why the two reference geometries require diffusion coefficients that differ almost threefold.

**Figure 9.**
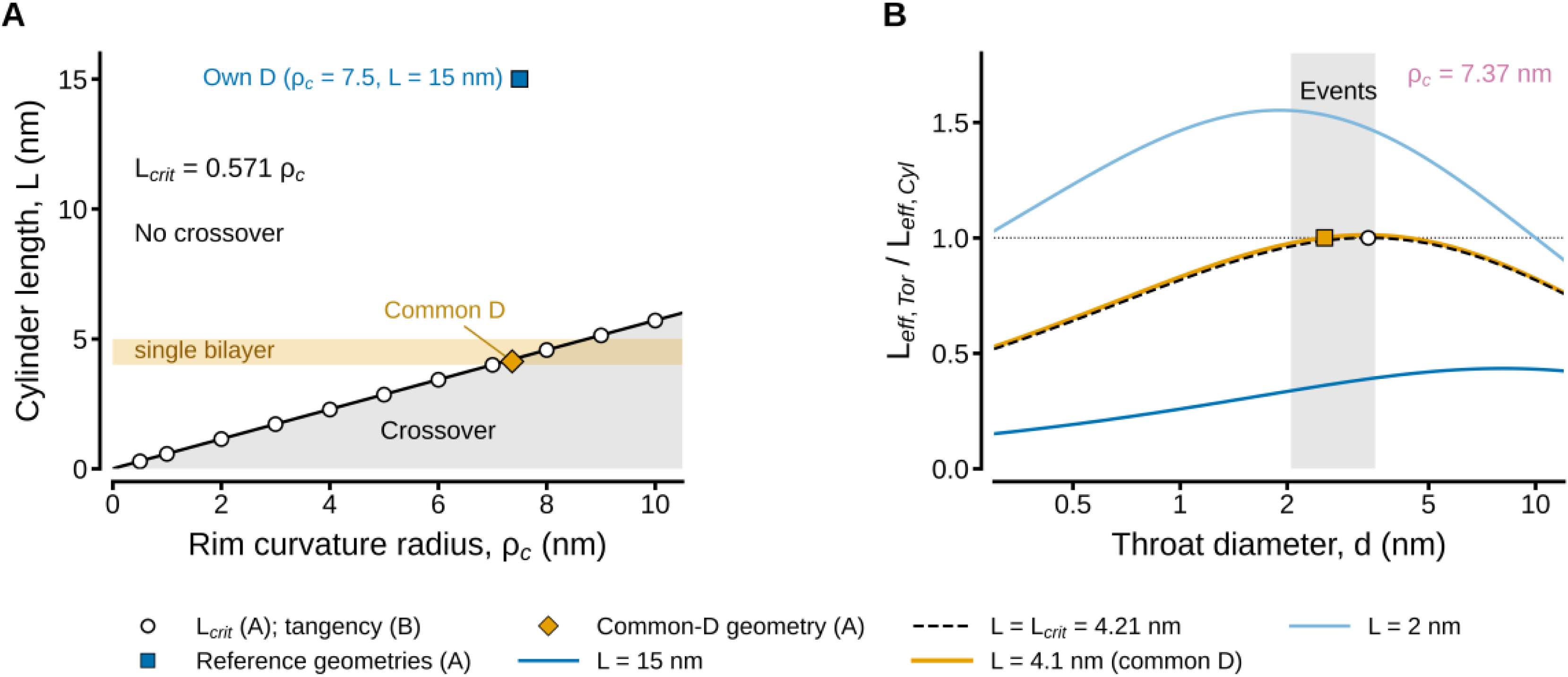
Matching of cylinder and toroidal resistance at equal. **D.** (A) Critical cylinder length, *L*_crit_, below which the resistances of a cylinder and a toroid with rim curvature radius *ρ_c_* coincide at some throat diameter (circles, *L*_crit_ = 0.571*ρ_c_*). Shaded gray area indicates the crossover region; orange band is the thickness of a single bilayer (4–5 nm). Square denotes reference geometries (*ρ_c_* = 7.5 nm, *L* = 15 nm); diamond denotes common-*D* geometry (*ρ_c_* = 7.37 nm,*L*_Cyl_ = 4.1 nm). (B) Ratio of toroidal to cylindrical effective length, L_eff,Tor_/L_eff,Cyl_, versus throat diameter for *ρ_c_* = 7.37 nm across cylinders of 15 nm, *L*_crit_ (4.21 nm; dashed line), 4.1 nm (common *D*), and 2 nm length. Gray band highlights peak event diameters (2.05–3.54 nm ); dotted line marks equal resistance; square indicates calibration diameter of the median event (2.55 nm); circle marks tangency at *L*_crit_ (*d* = 3.39 nm).

## Discussion

Our data favor a toroidal pore with fixed rim curvature as the geometry that releases serotonin during kiss-and-run exocytosis. The toroidal shape described all events with a single shape. Its kinetics were independent of precise rim curvature, its faster closure was robust to *D*, and it accommodated a common geometry-independent coefficient with almost no change in shape. A short cylinder of single-bilayer length, also fixed for all events, reproduced the amplitude range about as well, but its length depended strongly on the choice of *D*. Moreover, it narrows the kinetic signatures of a curved rim.

Four elements of the present approach are new: 1. The use of the geometry–*D* degeneracy as a test of physical admissibility; 2. A coefficient common to both geometries, the diffusional counterpart of the single resistivity used to size pores from conductance (Wu et al., 2017); 3. The exponent *γ* as a kinetic signature, independent of *D*, that identifies the rate-limiting structure; and 4. An event-specific quantal content, estimated by frozen pore extrapolation (De-Miguel, 2026), which prevents differences in vesicle content from being misinterpreted as differences in geometry. Previous reconstructions assumed a single geometry (Wu et al., 2017; Kapila and Nair, 2025) or treated geometry and *D* as irrelevant (Oleinick et al., 2013, 2017). A multiplicity of geometric and diffusional solutions has been noted (Kapila and Nair, 2025) but has not been used to discriminate between pore shapes.

### Comparing two geometries to identify a candidate pore

Because amperometry fixes only the ratio *L_eff_*/*D*, no single reconstruction can identify the pore. However, comparing two geometries under a common constraint narrows the choice. In our study, a candidate pore should meet four conditions: one fixed geometry reproduces the HFP-predicted peak diameter of every event; its geometric parameter remains physically admissible as *D* varies within plausible limits; its kinetic signature is robust to the choice of *D* ; and its parameters correspond to a known membrane structure. The toroid meets all four. The cylinder meets the first condition but not the second and third, since its length varies from 15 to 4.1 nm with *D* and its kinetic signature differs; it meets the fourth only at the short length, which matches a single bilayer (Chernomordik and Kozlov, 2008; Bao et al., 2016).

#### Determination of D

The diffusion coefficient of serotonin within a fusion pore remains unknown, although the diffusivity of serotonin associated with the dense-core vesicle matrix has been determined to be 2.1 × 10^−8^ cm^2^/s (Marszalek et al., 1997). This is the lower value used here as a reference limit, far from the *D* values we determined and well below the free-serotonin *D* value. In pores a few times wider than the solute, diffusion is hindered by steric exclusion at the entrance and by hydrodynamic drag at the walls. Therefore, *D* falls below its free-solution value by an amount that depends on the ratio of solute to pore size (Renkin, 1954; Dechadilok and Deen, 2006). To reach the pore, serotonin leaves the vesicle matrix via ion exchange, thereby lowering its effective mobility (Marszalek et al., 1997). Matrix proteins such as chromogranins bind stored amines and delay their exit after fusion (Borges et al., 2010), and fast transmitter discharge cannot be accounted for by free diffusion alone (Khanin et al., 1994). However, it is reasonable to suppose that once in the pore, serotonin flows according to a local *D* value. Pore sizes derived from conductance face the same uncertainty and assume the bulk resistivity of the solution for any geometry (Wu et al., 2017).

In model fittings, when *D* is left free, each geometry is calibrated to the same reference diameter using its own *D*, allowing *D* to absorb geometric differences. In this study, the values differed by almost threefold (7.7% and 21.1% of *D*_free_) for the same solute crossing pores of similar width. Neither coefficient can be transferred to the other geometry. The cylinder with *D*_Tor_ exceeded the HFP diameter ∼ 1.7-fold; the toroid with *D*_Cyl_ reached about half of it (Figure 1, A″–C″). These differences between reconstructions cannot be attributed to pore shape. A common *D* removes that degree of freedom and makes *D* a shared constant, so any remaining difference, and any geometric parameter that must change to absorb it, reports on the pore itself.

*D*_common_ describes diffusion of serotonin through the same aqueous lumen in different pores of similar width. There is no physical reason to expect serotonin to diffuse three times faster in a cylinder than in a toroid, as with geometry-specific coefficients. On the contrary, the *D* values absorb whatever each geometric factor gets wrong about the real pore, such as hindrance (Renkin, 1954; Dechadilok and Deen, 2006), entry effects (Hall, 1975), or wall shape.

Moreover, the 4.1 nm cylinder length under the common *D* coincides with the thickness of a single bilayer (Bao et al., 2016) and lies just below 0.57*ρ*_c_ ≈ 4.2 nm, the longest cylinder that can still match the resistance of the toroid (Figure 9); it therefore matches the thickness of a hemifusion diaphragm (Chernomordik and Kozlov, 2008). This possibility would not have appeared without the common-*D* strategy.

Conductance-based estimates of pore size use a single solution resistivity for any assumed geometry, for example, a 15-nm cylinder (Wu et al., 2017). A common *D* is the diffusional counterpart to that convention. Analyzing data both ways here yielded pore diameters comparable to conductance-based sizes (Breckenridge and Almers, 1987; He et al., 2006; Wu et al., 2017; Bao et al., 2018). Unlike those estimates, ours rest on the amperometric current alone, with no independent electrical measurement of the same pore. For conductance, the cylinder formula tested on ion channels of known structure predicts values about three times higher than measured. Such an approach gives a lower bound for the pore radius (Chang et al., 2017). The low *D* values obtained here relative to *D*_free_ may reflect a similar effect. Patch amperometry, which records the conductance of the fusion pore and the transmitter flux through it in the same vesicle (Albillos et al., 1997), would provide the independent measurement needed to separate geometry from *D*.

## The ***γ*** coefficient

Oleinick et al. (2017) analytically reconstructed pore radii from individual amperometric spikes, linking spike shapes to internal vesicle structure (Dannaoui et al., 2024). Like our approach, their model relies on the ratio of current to remaining intravesicular transmitter and an external calibration. Because flux is assumed to be limited by convergent matrix diffusion, the radius scales directly with this ratio, making pore geometry negligible. This corresponds to the access-limited boundary, *γ* = 1, in our framework. Applied to the median event resistance time-course, *γ* = 1 shortens the half-decay to 22.8 ms, versus 29.4 ms for toroidal and 32.9 ms for cylindrical geometries. This demonstrates that the rate-limiting step affects reconstructed kinetics more than pore shape. While this matrix-limited assumption holds for large dense-core vesicles expanding to ∼ 17 nm radii (Oleinick et al., 2017), it fails for sub-nanometer pores. For kiss-and-run pores (2–3 nm diameter), the pore itself limits flux, and its exponent depends on geometry. The diffusion correction factor reconciles geometries only at the calibration diameter, not across the full event.

### The short cylinder as a pore through a hemifusion diaphragm

The 4 nm length established for the cylinder matches the thickness of a single bilayer (Bao et al., 2016) and thus that of a hemifusion diaphragm formed by the distal leaflets after the proximal leaflets merge through a stalk (Chernomordik and Kozlov, 2003, 2008). A pore in such a diaphragm is a short channel through one bilayer (D’Agostino et al., 2016; Golani and Schwarz, 2023; Spencer et al., 2024). In simulations, these pores nucleate at the diaphragm rim and flicker for tens of nanoseconds (Spencer et al., 2024); a hydrophilic pore is preceded by a hydrophobic defect (Bennett et al., 2014; Akimov et al., 2017). The path from stalk to pore traverses low-energy intermediates (Kuzmin et al., 2001; Marrink and Mark, 2003; Knecht and Marrink, 2007), and high curvature lowers the opening barrier (Golani and Schwarz, 2023). Hemifused profiles have been imaged in chromaffin cells (Zhao et al., 2016), although they may reflect arrested fusion (Abbineni et al., 2018), and flickering pores form in hemagglutinin-mediated and protein-free fusion (Melikyan et al., 1993; Chanturiya et al., 1997). The short cylinder therefore describes a nascent rim pore that either collapses or grows into a neck on its way to the toroidal pore (Risselada et al., 2014).

### The toroid as the expanding pore

A pore that has grown into a lipid neck is a toroid, as described by theory and by images of hourglass-shaped pores (Razinkov et al., 1998; Wu et al., 2017). Wu et al. (2017) sized single pores with a 15 nm cylinder but modeled their energetics as a toroid using Helfrich bending energy (Helfrich, 1973). Expanding the nascent pore (radius ≈ 0.5 nm) threefold costs about 6 *k*_B_*T* with one SNARE but only 2 *k*_B_*T* with fifteen per face, because SNAREs crowding at the waist generate an entropic expansion force. Our reconstructions agree with this picture. One rim curvature describes all events; therefore, *γ* is insensitive to *ρ*_c_. A faster closure follows from a rim that does not scale with the throat. That is the defining property of a lipid neck. The 2.1–3.5 nm peaks lie close to the ∼ 3.4 nm boundary separating non-dilating from dilating pores (Wu et al., 2017), as expected for kiss-and-run.

### An integrated sequence

A sequence for a kiss-and-run event can now be proposed. SNARE zippering fuses the proximal leaflets into a hemifusion diaphragm without flux (Chernomordik and Kozlov, 2008; Zhao et al., 2016). A short cylinder then nucleates through the diaphragm, preferentially at its rim, with SNARE transmembrane domains at its edge (Bao et al., 2016; D’Agostino et al., 2016; Spencer et al., 2024; Melikyan et al., 1993; Chanturiya et al., 1997). The 4 nm cylinder spans the bilayer, over which syntaxin transmembrane segments may line the initial pore (Han et al., 2004). The pore grows into a toroid lined by SNARE complexes (Bao et al., 2016; Wu et al., 2017), and the toroid matches the lipidic neck into which it would dilate (Zhang and Jackson, 2010). Because ***γ*** depends only on wall shape, SNARE transmembrane domains at the waist are compatible with the toroidal signature (Bao et al., 2016; Wu et al., 2017; Dhara et al., 2020). The toroid occupies most of the rise and reaches 2.1–3.5 nm, sufficient for passage of 4–10 hydrated serotonin molecules. It does not dilate further (Wu et al., 2017; Bao et al., 2018; Trouillon and Ewing, 2014) and closes faster than a fixed-length channel because its rim does not scale with its throat, releasing only a fraction of the quantal content.

## Data availability

The amperometric events analyzed here are those of the companion paper (De-Miguel, 2026). The reconstructed time courses, parameter values and Python code used to generate the reconstructions and figures are deposited in a public repository (Zenodo; DOI to be added).

## Acknowledgments

I wish to acknowledge Mrs. Sara Cruz for the maintenance of the laboratory. This study was funded by a grant from the Human Frontiers Science Program Organization RGP0060/2019–102 to FFD-M, and by grant AG200823 from the Dirección General de Asuntos del Personal Académico, Universidad Nacional Autónoma de México, to FFD-M and Martha Takane.

## Abbreviations

5-HT: 5-hydroxytryptamine (serotonin)
Cyl: cylinder
HFP: hydrated flux packing
KR: kiss-and-run event identifier
Tor: toroid.

